# HMG Box-Containing Protein 1 (HBP1) Protects Against Pancreatic Injury in Acute Pancreatitis but Promotes Neoplastic Progression

**DOI:** 10.1101/2025.01.24.634817

**Authors:** Seung-Won Lee, Taelor Ekstrom, Elise C. Manalo, Shangyuan Ye, Mark Berry, Dmytro Grygoryev, Malwina Szczepaniak, Jinho Lee, Carlos Origel Marmolejo, Syber Haverlack, Juyoung Lee, Vidhi M. Shah, Dove Keith, John L. Muschler, Koei Chin, Rosalie C Sears, Stuart P. Weisberg, Terry Morgan, Jungsun Kim

## Abstract

**Background & Aims:** Pancreatitis is an inflammatory disease of the exocrine pancreas and a known risk factor for pancreatic ductal adenocarcinoma (PDAC). Previously, we identified HMG-box transcription factor 1 (HBP1) as a potential master transcription factor (TF) in the early progression of PDAC, with its expression associated with poor patient survival, underscoring its significance in pancreatic disease. However, the functional role of HBP1 in the onset and progression of acute pancreatitis (AP) remains unknown.

**Methods:** We examined HBP1 expression in human pancreatitis samples and a cerulein-induced AP mouse model. Pancreatic-specific conditional HBP1 knockout mice, with or without an oncogenic *Kras* mutation, were generated and compared to their littermate controls. Spatial transcriptomics and multiplexed protein assays, histological analysis, and immunostaining were utilized to characterize pathological changes. Findings from mouse models were validated using inducible HBP1-overexpressing human pancreatic ductal epithelial cells.

**Results:** HBP1 was upregulated in pancreatic exocrine cells in human chronic pancreatitis and mouse acute pancreatitis, with its expression in human chronic pancreatitis correlating with cancer presence. Pancreatic HBP1 ablation disrupted acinar homeostasis by impairing autophagic flux and exacerbating inflammation following injury. In the presence of oncogenic *KRAS*, HBP1 ablation delayed the formation of pancreatic intraepithelial neoplasia (PanIN), the precursor to PDAC, and slowed its progression to higher-grade lesions.

**Conclusions:** HBP1 upregulation in pancreatitis mitigates pancreatic inflammatory injury; however, in the presence of oncogenic *KRAS*, it facilitates PanIN progression. Thus, HBP1 serves as a critical regulator in both pancreatitis and early pancreatic neoplasia, representing a potential therapeutic target for intervening pancreatitis and PanIN progression.

## INTRODUCTION

Acute pancreatitis (AP) is an inflammatory disease of the exocrine pancreas caused by acinar cell damage, leading to autodigestion, inflammation, and necrosis^1–4^. While most AP cases are mild to moderate and resolve within days or weeks, approximately 20% of AP cases develop severe disease that can be lethal^1–4^. Recurrent AP increases the risk of chronic pancreatitis (CP), which is predisposed to developing pancreatic neoplastic lesions^1–4^. However, a single severe AP episode does not significantly increase the risk of recurrence of AP but repeated mild AP episodes elevate this risk. Thus, the relationship between severe AP and the onset of CP is not linear. Therefore, a deeper understanding of factors and processes regulating AP severity is critical for predicting outcomes and preventing its recurrence and progression to CP and neoplastic lesions^1–4^.

Pathologically, mild AP is characterized by interstitial edema within the exocrine compartment, accompanied by a focal increase in immune cells but without acinar dropout. Moderately severe AP is defined by focal acinar dropout with immune cell infiltration, whereas severe AP is marked by diffuse (multifocal) acinar dropout, intense immune infiltration, or features of CP^4, 5^. Pathophysiologically, AP is broadly classified into two main types^4^: (i) mild interstitial edematous pancreatitis, which occurs in most patients and typically resolves within a week. This form is characterized by homogeneous pancreatic enlargement due to inflammatory edema, mild peripancreatic fat inflammation, and an absence of tissue necrosis. (ii) Severe necrotizing pancreatitis, observed in 5–10% of cases, involves inflammation associated with both parenchymal and peripancreatic necrosis^4^.

Several factors can induce and exacerbate pancreatitis. Premature activation of pancreatic enzymes leads to pancreatic injury, including acinar cell death, and triggers inflammation. A persistent inflammatory response further aggravates disease severity^6–8^. Autophagy plays a crucial role in maintaining pancreatic acinar homeostasis under normal conditions^6^. Autophagic flux encompasses the entire process of autophagy, from autophagosome formation to its fusion with lysosomes and subsequent degradation. Disruptions in autophagic flux compromise acinar homeostasis, leading to aberrant cell death. Impaired autophagic flux is a hallmark of AP, as evidenced by the accumulation of large autophagic vacuoles marked by p62 and LC3 in acinar cells^6^.

For inducing AP experimentally, repeated intraperitoneal injections of cerulein are the most widely used and reproducible method^9, 10^. Cerulein stimulates acinar cells to produce excessive digestive enzymes as well as inappropriate secretion of pancreatic enzymes and production of reactive oxygen species that induce inflammation and acinar cell death^9, 10^. Similar to human AP, cerulein treatment induces histological changes, such as interstitial edema, leukocyte infiltration, and acinar cell dropout, that peak at 3-6 hours post-treatment. These changes begin to resolve by 24 hours, with the pancreas appearing morphologically normal after one week^11^. The presence of an oncogenic KRAS mutation, particularly with prolonged cerulein treatment, can induce CP-like features, including persistent acinar atrophy, inflammation, and stabilized acinar-to-ductal metaplasia (ADM). This progression can further lead to pancreatic intraepithelial neoplasia (PanIN), the most common precursor to pancreatic ductal adenocarcinoma (PDAC)^11–14^.

HMG-box transcription factor 1 (HBP1) regulates the G1-to-S cell cycle progression through binding RB^15, 16^. Upregulated HBP1 induces senescence and promotes DNA-break repair upon oxidative stress or oncogenic mutations^17–19^. Thus, HBP1 has traditionally been considered as a tumor suppressor. Yet, recent studies suggest that HBP1 may also function as a tumor promoter^20, 21^ and that HBP1 serves as a hub across multiple signal pathways through various post-translational modifications^22^. The role of HBP1, therefore, may be tissue- and context-dependent. We previously identified HBP1 is normally expressed only in islet cells of the adult pancreas but becomes upregulated in exocrine cells in PDAC and CP adjacent to PDAC in human^23^. Furthermore, we demonstrated that HBP1 functions as a potential master transcription factor (TF) in the early progression of PDAC, with its expression correlating with poor patient survival^23^. Despite these findings, the functional role of HBP1 in pancreatitis and the initiation of PanINs has not been reported. Herein, we investigated the role of HBP1 in AP and the initiation and progression of PanINs.

## MATERIALS and METHODS

### Mouse Model

All animal procedures were approved by the OHSU IACUC (IP00004592). *p48^cre^* (#023329) and *LSL-K-ras^G12D/+^* (#008179) mice (Jackson Laboratories) were backcrossed onto the C57Bl/6J background. *Hbp1^flox/flox^* mice (C57BL/6J, ICR) were derived from cryopreserved embryos, courtesy of Dr. Ohtsuka’s lab^24^. These mice were backcrossed for at least five generations with p48^Cre^ (C57BL/6J) mice (Fig. S2A). Genotypes were confirmed using PCR or real-time PCR (Transnetyx, Table S1). Pancreatitis was induced in 8–10-week-old mice via seven hourly intraperitoneal injections of 50 µg/kg cerulein (Sigma)^25^, and pancreata were collected 1, 2, and 7 days after the last cerulein injection.

To generate HBP1-null KCH (*p48^Cre^*; *LSL-Kras^G12D^*; *Hbp1^flox/flox^*) and HBP1-competent KCH (*p48^Cr^*^e^; *LSL-Kras^G12D^*; *Hbp1^wt/flox^*) mice, *Hbp1^flox/flox^* mice were crossed with *p48^Cre^; LSL-Kras^G12D^* (KC) mice. Pancreata were collected at 10, 20, and 40 weeks to assess i) body and pancreatic weights, ii) histology analysis, and iii) tumor burden in the liver, lung, and local lymph nodes.

### PanIN and PDAC Histopathology

A board-certified pathologist (TKM), blinded to genotype, assessed PanIN grades from H&E images^26^. Normal ducts display uniform cuboidal epithelium without mucinous changes or nuclear atypia. PanIN1 has a flat architecture, minimal budding, abundant mucin, and small oval nuclei. PanIN2 retains mucin but adopts a papillary structure with mild nuclear atypia. PanIN3 (carcinoma in situ) is characterized by pleomorphic nuclei, increased nuclear-to-cytoplasmic ratio, cribriforming, epithelial budding, luminal necrosis, and loss of polarity.

### Human Pancreatitis Tissue Microarray (TMA)

De-identified human pancreatitis tissues were obtained through the Oregon Pancreas Tissue Registry (IRB00003609) with informed consent. The OHSU IRB approved all protocols, following relevant guidelines. Chronic pancreatitis tissues were collected via fine-needle aspiration (FNA) or Whipple surgery from patients with intraductal papillary mucinous neoplasm (IPMN). Tumor-associated pancreatitis tissues were obtained from PDAC/Intraductal papillary mucinous carcinoma (IPMC) margin resections. The TMA includes 170 cores from 65 subjects (49 chronic pancreatitis, 16 controls), with each patient contributing 2–6 replicate cores. A total of 311 tissue regions were classified as pancreatic (Acini: n=89, Ducts: n=77, Stroma: n=132) or control tissues (Spleen: n=5, Placenta: n=4, Tonsil: n=4).

### Nanostring Digital Spatial Profiling (DSP) GeoMx Platform

Mouse immune profiling, activation, cell typing, and immune-oncology target modules were used^27^. Mouse protein panels were incubated on pancreata from four HBP1 WT and four CKO mice on days 1 and 2 post-cerulein treatment using PanCK (epithelial cells) and CD45 (immune markers). Ninety-six regions of interest (ROI) with pancreatitis features were selected for DSP analysis. UV-cleaved DNA oligos were hybridized with fluorescent codesets for molecule counting. nCounter data were normalized via geometric mean, and signal-to-noise ratio (SNR) per target protein was computed with IgG controls. Twenty-seven immune markers with log₂(SNR) > 1.6 were selected for downstream analysis. Geomean-normalized data were used for clustering and statistical analysis.

### 10x Visium Spatial Transcriptome

Spatial transcriptomic analysis was performed on pancreata from three HBP1 WT and three HBP1 CKO mice on days 1 and 2 post-cerulein treatment. Pancreatitis areas were mounted on 10x Genomics Visium slides, with 5000 barcoded 55 µm spots capturing 1–10 cells per spot, each containing millions of probes for 20,551 mouse genes via CytAssist^28^. Sequencing (25k read pairs/spot) generated FASTQ files, which were mapped to H&E images and feature-barcode matrices using Spaceranger. Seurat was used for data processing, analysis, and visualization^29^, while Harmony corrected technical variations across slides^30^.

### Statistical Analysis for Pancreatitis TMA

HBP1 expression in TMAs, assessed by IF staining, was quantified using the H-score (0 = negative, 1 = weak, 2 = moderate, 3 = strong), and the percentage of cells at each intensity was calculated per cell type using QuPath. The final H-score per cell type was obtained by multiplying the score by the percent of stained cells (0–100%), then dividing by 100, yielding a 0–3 range. Since each patient sample contained mixed cell types, combined scores were computed per patient. Logistic regression assessed the association between HBP1 expression (individual and combined cell types) and pancreatic tumors (PDAC+IPMC). Scores of each cell type are used as predictors, and the presence of a tumor is used as the outcome variable. ROC curves evaluated the model’s predictive power.

### General Statistical Analysis

Animals were age- and sex-matched for each experiment. Immunostaining and western blot quantifications were performed using QuPath and ImageJ, respectively. GraphPad Prism assessed data distribution. Welch’s t-test and Mann-Whitney test were used for pairwise comparisons of normally and non-normally distributed data, respectively. A two-part model was applied for Fig. 2G due to a high proportion of zero values. Significance threshold: p < 0.05. A hypergeometric test assessed randomness, while the Benjamini-Hochberg procedure controlled the false discovery rate (FDR). Spearman’s rank correlation (ρ) evaluated molecular signature reproducibility.

**Tissue immunofluorescence (IF), immunohistochemistry (IHC), H&E staining, and quantitative reverse transcription-PCR (qRT-PCR)** were performed as previously described^23^.

## RESULTS

### The expression of HBP1 is increased in pancreatitis and is associated with the pancreatitis progression

To investigate the role of HBP1 in pancreatitis and PanINs, we ascertained HBP1 expression in mouse cerulein-induced AP and human pancreatitis tissues using a verified HBP1 antibody (Fig. S1A-B). In mice, HBP1 was scarce in normal pancreatic exocrine cells (Fig. 1A-B) but was widely increased in damaged pancreatic acinar cells after the cerulein treatment, reaching the maximum at 48 hours after the final cerulein treatment (Fig. 1C-D, S1C-G). Concurrent with the recovery from injury, the HBP1 protein disappeared by day 7 after cerulein treatment (Fig. 1E, S1H-I). These results indicated that the HBP1 expression increased during cerulein treatment-induced pancreatitis and correlated with its kinetics.

**Figure 1.**
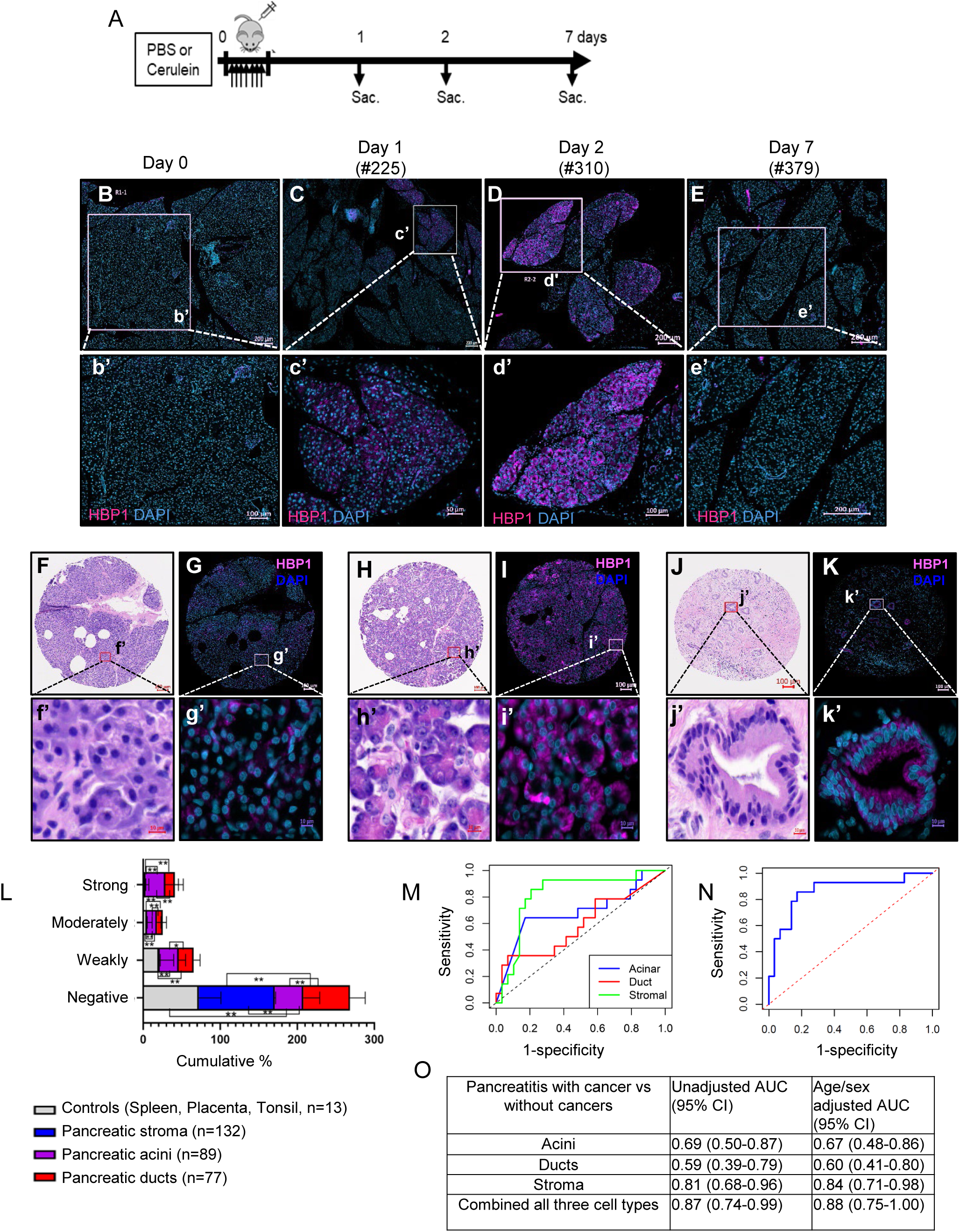
The expression of HBP1 shifts from pancreatic islets to acinar cells in pancreatitis. A. Schematic diagram of cerulean-induced AP model. B-E. Representative images of HBP1 immunostaining in the pancreata of one set of mice on day 0 (B, inset b’), 1 (C, inset c’), 2 (D, inset d’), and 7 (E, inset e’). n=3 each time point, images from the rest of the mice are shown in Fig. S1. F-K. Representative H&E images (F, H, J) and immunofluorescent images of HBP1 expression (G, I, K) on normal-looking acinar (F, G), injured acinar cells (H, I), and PanINs cells (J, K) from patients with pancreatitis. Scale bars represent 100 µm (top images) or 10 µm (bottom images). L. Summary of HBP1 expression in human pancreatic tissues (311 samples from 49 patients with pancreatitis and 16 normal subjects). HBP1 expression is significantly higher in injured acinar cells and PanINs compared to stromal cells or normal controls (*p < 0.05, **p < 0.001; unpaired Welch’s t-test, Table S2D). M-O. ROC curves and AUC values of HBP1 expression for each cell type (M, O) and for the combined cell types (N, O) in pancreatitis patient tissues, which were analyzed against the presence of pancreatitis with cancer.

In human tissues, HBP1 was barely or only weakly expressed in normal acinar tissues, stroma, and other normal tissues (Fig. 1F-G, L) but was significantly expressed in injured acinar and ductal cells in pancreatitis tissues (Fig. 1H-L, Table S2A-D). We investigated the association between HBP1 levels in pancreatitis tissues and invasive tumors using a logistic regression model (see Methods). HBP1 expression in fibroblast cells was significantly inversely correlated with tumor presence, suggesting a protective role. In acinar and ductal cells, HBP1 showed negative and positive associations with tumor presence, respectively, though not statistically significant, possibly due to the complex roles of these compartments or limited sample sizes (Fig. S1J-K). These findings highlight the cell type-specific role of HBP1 in pancreatic disease progression.

Next, we evaluated whether HBP1 expression in individual or combined cell types could differentiate pancreatitis patients with invasive tumors using receiver operating characteristic (ROC) curve analysis. The area under the ROC curve (AUC) values were 0.67 (95% CI: 0.48-0.86) for acinar, 0.60 (95% CI: 0.41-0.80) for ductal, and 0.84 (95% CI: 0.71-0.98) for stromal cells. Combining all three cell types improved performance (AUC = 0.88, 95% CI: 0.75-1.00) compared to individual cell types. Gender and age were included as variables in the model but were not significant (p = 0.55 and 0.19, respectively) (Fig. 1M-O).

In summary, HBP1 is ectopically overexpressed in cerulein-induced mouse AP and human CP tissues, and its expression in human CP may be linked to the presence of invasive tumors within pancreatitis.

### Loss of Hbp1 in the pancreas shows a grossly normal phenotype but exacerbates the AP severity

To investigate the pathophysiological functions of HBP1 during pancreatic inflammation and regeneration, we generated pancreatic-specific Hbp1 conditional knockout (“CKO”) mice by crossing *Hbp1^flox/flox^* mice^24^ with p48^Cre^ mice, in which *p48 (Ptf1a)*^Cre^ recombinase is expressed specifically in pancreatic progenitor and adult acinar cells^31^ (HBP1 CKO mice: *p48^Cre^; Hbp1^flox/flox^*) (Fig. S2A-C). *Hbp1^flox/flox^* mice were used as “littermate wild type (WT) controls”. Both sexes of HBP1 CKO mice developed normally and were born at expected Mendelian ratios with normal histology indistinguishable from control mice. We verified that HBP1 CKO mice had decreased expression of HBP1 in the pancreatic islet compartment (Fig. S2D-G).

To assess the *in vivo* function of HBP1 in pancreatic injury, we induced cerulein-mediated AP in HBP1 CKO and littermate WT control mice (Fig. 2A). Cerulein treatments had minimal impact on the body weight in both groups yet increased normalized pancreatic weights compared to PBS treatment one-day after, reflecting edematous AP (Fig. 2B-C, Table S3). The normalized pancreatic weights of HBP1 WT and CKO mice were not different on day 1. However, those of HBP1 CKO mice were significantly lower than those of HBP1 WT control mice by day 2, indicating parenchymal destruction/atrophy that was often associated with severe pancreatitis (Fig. 2B-C).

**Figure 2.**
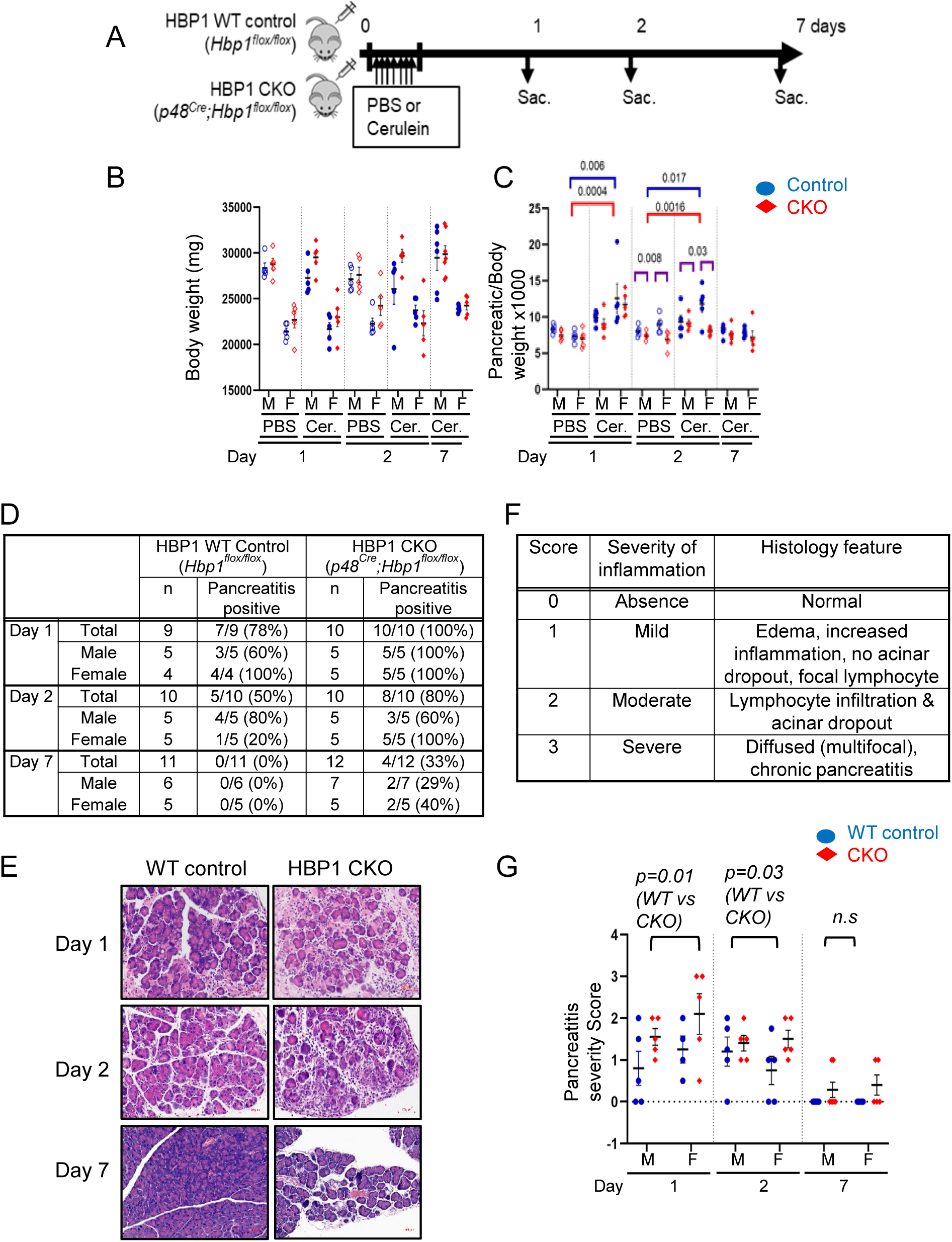
Hbp1 ablation exaggerates AP phenotype in the cerulean-induced AP model. A. Diagram of cerulein-induced AP model. Cerulein was injected every hour for seven injections. Mice were sacrificed 1, 2, and 7 days after the last injection and their pancreata were investigated. B-C. Body weight (B) and pancreatic weight normalized with body weight (C) of mice of indicated sexes at each time point. There is no significant difference in body weight between female and male mice (B). Compared to PBS controls, normalized pancreatic weight is significantly higher in cerulein-treated both HBP1 WT (p-values are labeled in blue) and HBP1 CKO (p-values are labeled in red) mice on day 1 and day 2. Normalized body weight significantly decreased in HBP1 CKO compared to WT mice in either PBS or cerulein-treated mice on day 2 (p-values are labeled in purple). The unadjusted p-values are calculated based on the two-sample t-test without the equal variance assumption. Linear models include group indicators and sex indicators as covariates that are fitted to the outcome of interest (weight, pan weight, or norm pan weight). The p-values corresponding to the coefficient of the group indicator are reported as the adjusted p-values. D. Binary scoring table of the presence of pancreatitis. The presence of pancreatitis was determined by the pathologist based on H&E images in an unblinded manner. E. Representative H&E images of pancreatitis in HBP1 WT control and HBP1 CKO mice. Scale bars indicate 50 µm. F. Pancreatitis severity scoring table, based on H&E images. G. The severity of pancreatitis in HBP1 WT control and CKO mice was determined on days 1, 2, and 7 after cerulein induction. Statistical analysis was made using two-part fitted models (p=0.01 and 0.03 for days 1 and 2, respectively, n=10-12 per group at each time point).

AP, determined by a pathologist based in a blinded manner on H&E staining images^4^, occur at a higher frequency in HBP1 CKO mice than in control mice upon cerulein treatments (Fig. 2D-E). Because we did not observe any apparent associations of sex and age with the pancreatic injury, we did not include sex and age as adjustment variables in subsequent modeling analyses.

Next, we assessed the severity of AP in the pancreata of HBP1 WT and CKO mice based on H&E staining images^4^ (Fig. 2F). The exocrine compartments of cerulein-treated HBP1 WT mice displayed mild injury with degranulated acinar cells and interstitial edema. In contrast, cerulein-treated HBP1 CKO mice showed diffuse and more extensive acinar dropout, indicating a significantly increased severity of AP (Fig. 2E-G). Since we used a cerulein treatment condition for mild AP^25^, we did not observe a significant increase in the ADM population in either HBP1 WT or CKO mice (Fig. S2H). Subsequently, as expected, the pancreas in both HBP1 control and CKO mice was repaired with regenerated acinar cells, and its overall morphology was indistinguishable from that of PBS-treated animals seven days after cerulein treatments, although focal areas exhibited sustained mild AP in a subset of CKO mice (Fig. 2E-G).

Collectively, these data indicate that HBP1 protects the pancreata from initial pancreatic injury and fibrosis but is dispensable for pancreatic metaplasia and regeneration.

### Loss of Hbp1 increases inflammation that is largely driven by macrophages in AP

As HBP1 ablation increased pancreatic injury, we sought to identify leukocytes that differentially infiltrated the pancreata of HBP1 CKO mice after cerulein treatment. Because contaminations by leukocytes from lymph nodes during tissue dissociation could exaggerate the extent of leukocytes infiltrating the pancreas when using flow cytometry, we used the DSP GeoMx platform with mouse immune cell panels^27^ to spatially resolve cellular components (Fig. 3A). Pancreatic cores from HBP1 WT and CKO mice exhibited distinct clustering (Fig. 3B). Consistent with a prior report^32^, HBP1 CKO pancreata showed increased Ki67 and GAPDH (Fig. 3C). Importantly, HBP1 CKO pancreata had significantly increased myofibroblast (Fibronectin, SMA) and myeloid lineage (CD11b, F8/40, CD44 and VISTA) populations as well as total immune cells (CD45) but had decreased T-cell tolerance signals (OX40L, CD4 and FoxP3) on days 1 and 2 after cerulein treatment (Fig. 3D-E, S3A-C).

**Figure 3.**
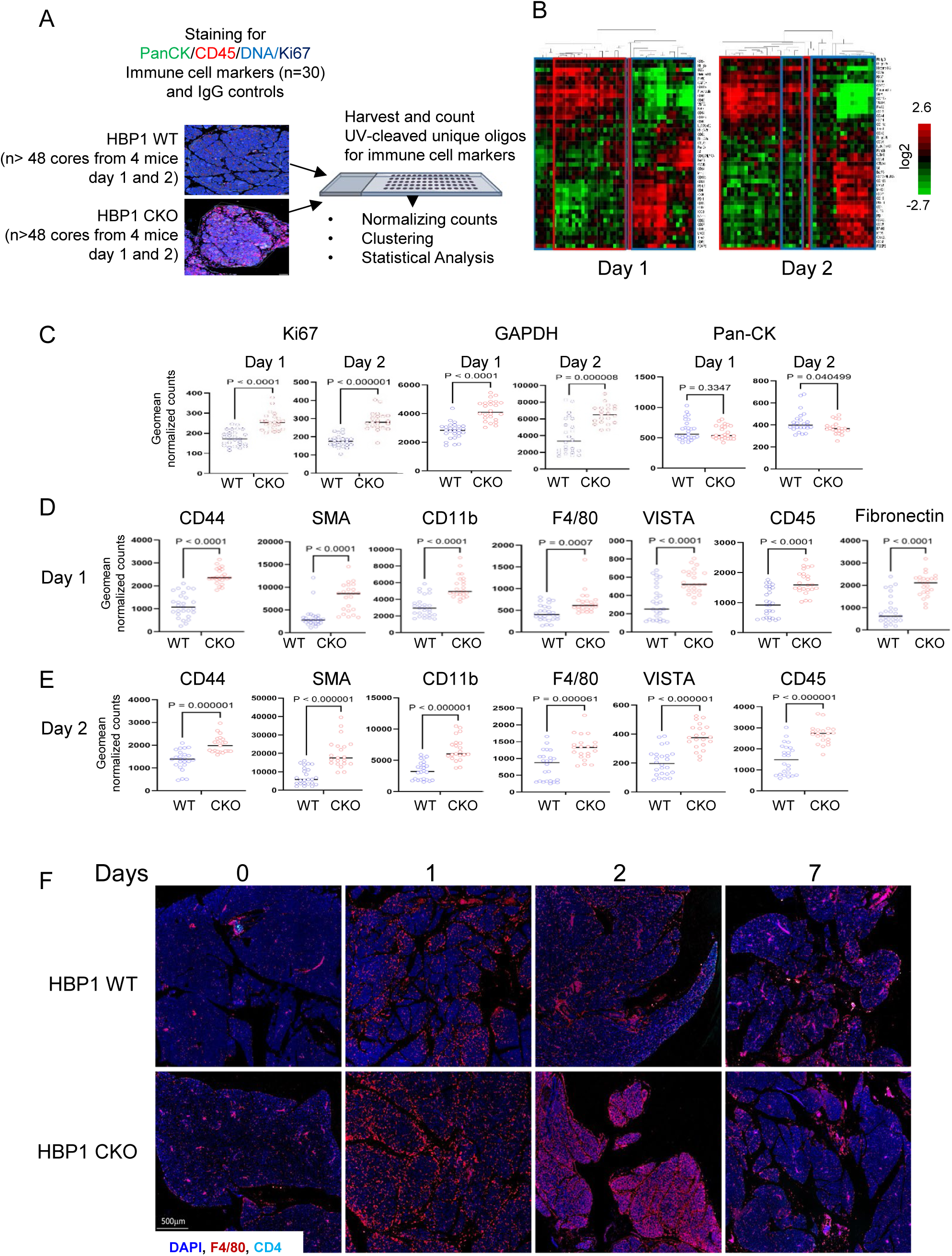
HBP1 ablation increases myofibroblasts and myeloid lineage cells in AP. A. Schematic diagram of DSP. A total of 24 pancreatitis cores from two mice per group in each time point were applied for the DSP mouse panels, including immune cell profiling core, immune activation status, immune cell typing and immune-oncology drug target protein modules^27^. B. Unsupervised hierarchical heatmap clustering analysis. Core samples and protein targets were clustered in the 2-way arrays using the Euclidean method. Blue brackets indicate the pancreata from HBP1 WT mice, and red brackets indicate the pancreas from HBP1 CKO mice. C. The expression of marker proteins for proliferation (Ki67), metabolism (GAPDH), and of Pan-CK proteins. D-E. Significantly increased immune cell markers in HBP1 CKO mice compared to WT mice on day 1 (D) and 2 (E) post-cerulein. F. Representative images of immunostaining for F4/80^+^ macrophages and CD4^+^ cells.

To better spatially resolve leukocytes in pancreatic inflammation, we performed IF with markers of T cells (CD3, CD4) and macrophages (F4/80). The infiltration of F4/80^+^ macrophages into the pancreatic parenchyma was markedly higher in HBP1 CKO compared to control mice at days 1 and 2 after cerulein treatment. In contrast, there was no detectable T cell infiltration into the pancreatic parenchyma during this period in these mice (Fig. 3F). In both groups, we observed a marked accumulation of CD4^+^ T cells in the center of peripancreatic lymph nodes with a dense rim of F4/80^+^ macrophages around the periphery of the lymph nodes on days 1 and 2 after cerulein treatment (Fig. S3D). These results show that the increased inflammation in pancreatic parenchyma of HBP1 CKO mice during cerulein-induced pancreatitis is largely macrophage-driven, whereas CD4^+^ T cells and macrophages accumulate in the peripancreatic lymph nodes similarly in the HBP1 WT and CKO mice.

### Loss of Hbp1 enhances immune response gene programs but impairs pancreatic protective gene programs upon stress in AP

To understand how HBP1 ablation impaired acinar homeostasis and triggered the infiltration of macrophages, we characterized the spatial transcriptomics of the pancreas of HBP1 WT and CKO mice collected on days 1 and 2 after the cerulein treatment using the 10x Genomics Visium system (Fig. 4A, n=3 each group, see methods). We obtained the median number of genes per spot between 500 and 5000 and the median number of UMI counts per spot between 2000 and 10000 (Fig. S4A, Table S4). After removing lymph node tissues, we integrated transcriptomes from all samples using Harmony^30^ to reduce the technical variations among tissue sections (Fig. 4A). The cell cycle did not affect the distribution of transcriptomes among samples, and the Spearman correlation coefficient among samples was above 0.9, ensuring further analysis (Fig. S4B-C).

**Figure 4.**
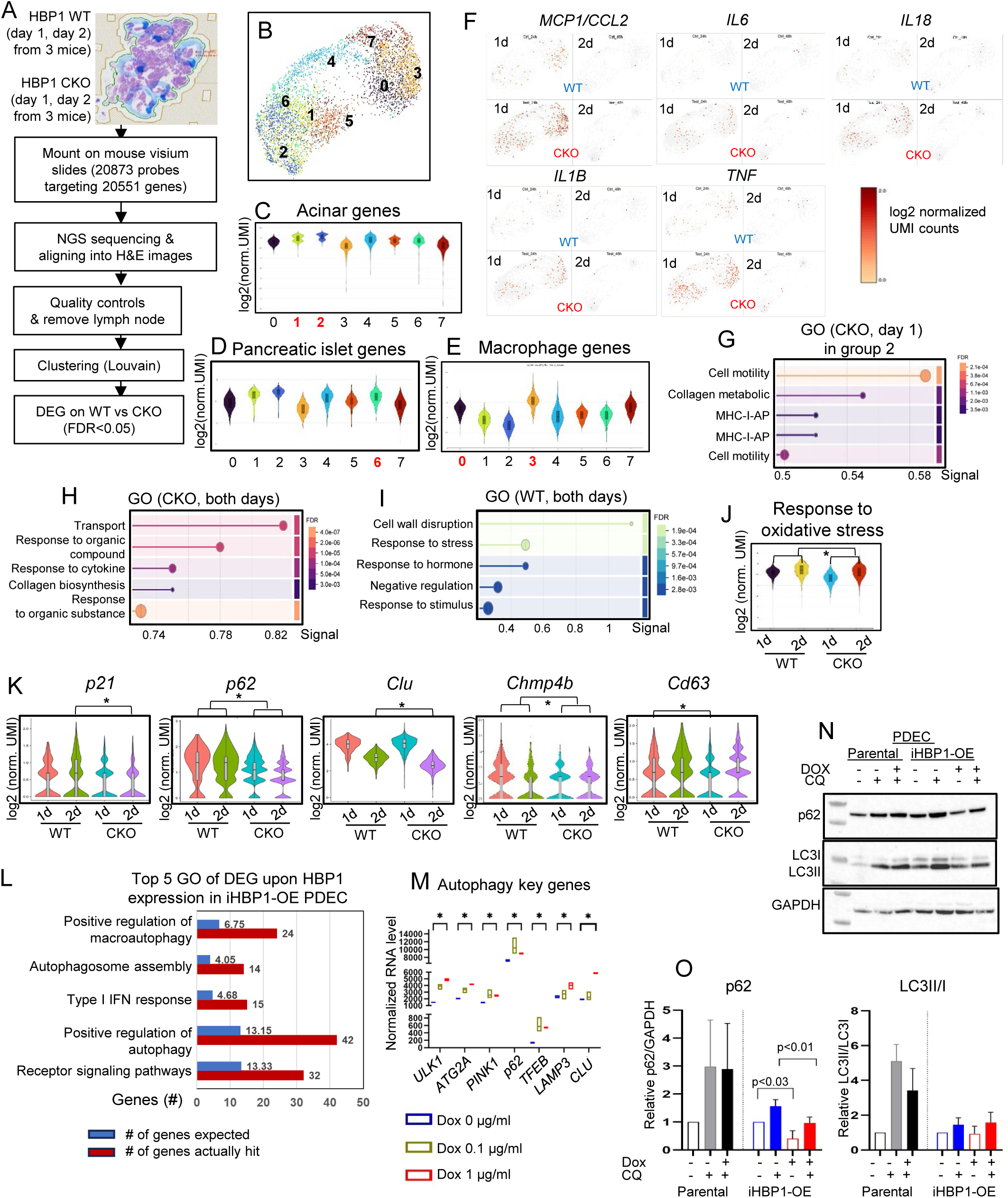
Hbp1 ablation enhances immune response gene programs and impairs cellular response to stress in the cerulean-induced AP model. A. Schematic diagram of Visium spatial transcriptomic analysis of pancreata derived from HBP1 WT and CKO mice on days 1 and 2 after cerulein treatments (n=3, each group). B. Uniform Manifold Approximation and Projection (UMAP) displayed eight clusters of integrated samples. C-E. The expression of pancreatic acinar (C), islet cells (D), and macrophage (E) marker genes. The X-axis indicates each cluster. The Y-axis indicates log2 transformed normalized UMI counts per spot. F. Mapping of proinflammatory genes in the UMAP. HBP1 WT or CKO pancreata are displayed in UMI. Each dot represents each spot expressing cytokine genes, displayed in log2-normalized UMI counts. G-I. The top five GO terms were identified for upregulated genes in group 2 cells on day 1 (G), genes upregulated across all groups at both time points (H), and genes downregulated across all groups at both time points (I), in HBP1 CKO compared to WT pancreata. The signal on the X-axis is defined as a weighted harmonic mean of the observed/expected ratio and −log (FDR). J-K. The expression of genes involved in the cellular response to oxidative stress (J) and autophagy pathways (K) was significantly downregulated in HBP1 CKO compared to WT pancreata (*FDR < 0.05). L. Top5 GO of DEG upon HBP1 expression in iHBP1 OE PDEC (FDR<0.05). M. Expression of autophagy genes among DEG increased as HBP1 levels increased (*, FDR<0.05). N-O. Western blot analysis of autophagy markers p62 and LC3 in iHBP1-OE PDEC cells treated with doxycycline (1 µg/ml, 24 hrs) to induce HBP1 overexpression, followed by chloroquine (CQ, 12 µM, 24 hrs) treatment (N-representative images). Quantification of p62 levels normalized to GAPDH and LC3-II/LC3-I ratio was performed. Band intensities were measured using ImageJ. Statistical analysis was performed using Welch’s t-test (n=4 for p62, n=2 for LC3) (O).

We clustered spatial spots into eight groups based on their spatial relationships and gene expression patterns within the tissue using the Louvain algorithm (Fig. 4A-B, S4D), with different groups representing distinct cell types. Group 0: neutrophils and macrophages; Groups 1 and 2: pancreatic exocrine cells; Group 3: myofibroblast and myeloid cells; Group 6: pancreatic islet cells; Group 7: collagen expressing fibroblast cells (Fig. 4B-E, S4E-H). Although cells from HBP1 WT and CKO pancreata were distributed across all clusters, those from CKO pancreata were slightly enriched in groups 0, 3, and 7, reflecting the severity of pancreatitis in CKO mice (Fig. S4I). The expression of genes encoding proinflammatory cytokines crucial for recruiting monocytes in HBP1 CKO mice was higher in cells from CKO pancreata, supporting that HBP1 ablation increased macrophage infiltration by regulating proinflammatory cytokine genes (Fig. 4F). Regenerating Gene (Reg) family proteins are crucial protective factors associated with pancreatitis to aid the recovery of pancreatic acinar cells by exerting anti-apoptotic and anti-inflammatory effects in response to oxidative stress in pancreatitis^33, 34^. Indeed, the expression of Reg genes was significantly lower in HBP1 CKO compared to WT pancreata post-cerulein treatment (Fig. S4J), confirming that the spatial transcripts reflected the pathophysiological events.

To understand gene programs impacted by HBP1 CKO in the pancreata, we characterized differentially regulated genes (DEG) between HBP1 WT and CKO pancreata in group 2 cells, which are mainly pancreatic acinar cells (Fig. 4B-C, S4E). Consistent with the main function of HBP1 as a transcriptional repressor, most DEGs were upregulated in HBP1 CKO pancreata (Fig. S4K). Programs related to the immune response were significantly increased in HBP1 CKO pancreatic group 2 cells (Fig. 4G). In contrast, genes encoding digestive enzymes, such as lipases (Pnliprp2 and Pnlip) and SPINK1/SPINK3 protease inhibitors, and molecules related to anti-inflammation (Reg3d, Tff2) and metabolism (Ggh, Nucb2) were among a small number of genes downregulated in CKO pancreata, indicating that HBP1 ablation reduced pancreatic protection against injury (Fig. S4L).

To gain insight into the effect of HBP1 ablation in the entire pancreas, we analyzed DEG in HBP1 CKO compared to WT pancreas across all cell groups. Reflecting the severity of pancreatitis, Spink1 was significantly downregulated in HBP1 CKO compared to WT pancreata, as expected (Fig. S4M).

Gene ontology (GO) of DEG indicated that, compared to WT pancreata, intracellular transport and cytokine response were significantly increased whereas pathways associated with response to stress, including oxidative stress, were significantly decreased in HBP1 CKO pancreata (Fig. 4H-J). Events such as apoptosis, senescence, and autophagy occur in response to oxidative stress^35–38^. Apoptotic pathways were decreased in both HBP1 WT and KO pancreata on day 2 after cerulein treatment (Fig. S4N-O), and apoptotic cells were minimally impacted by HBP1 ablation (Fig. 5A-D). HBP1 increases senescence^17, 18^. Interestingly, HBP1 ablation significantly downregulated autophagic flux genes, including *p21, p62,* and *Clu* (autophagosome formation) and *Chmp4b* and *Cd63* (degradation and secretion)^39–41^ (Fig. 4K).

**Figure 5.**
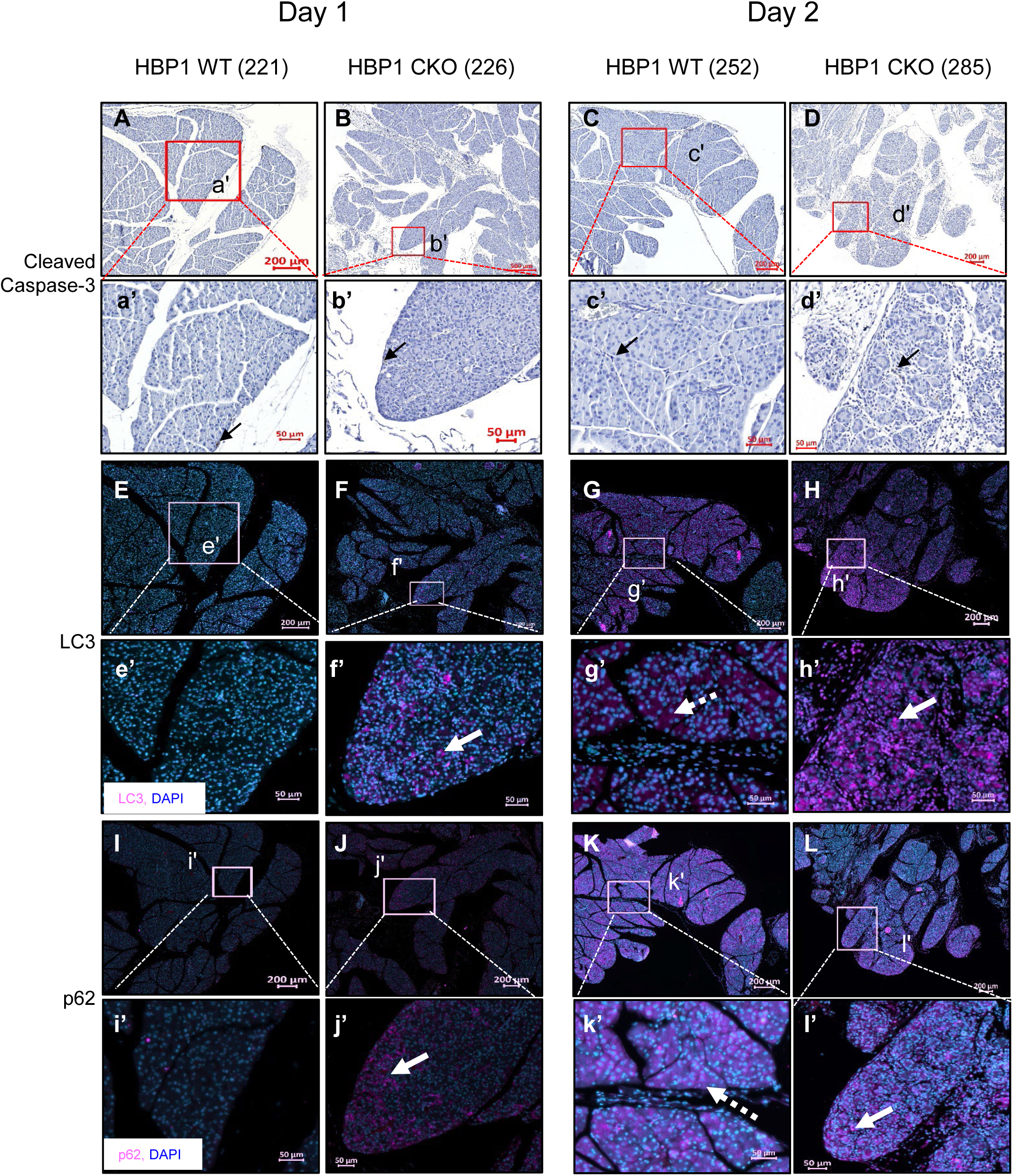
Representative images of immune staining for cleaved caspase-3 (A-D, arrow; positive signals) and IF for autophagosome marker proteins LC3 (E-H) and p62 (I-L) in pancreata from HBP1 WT and CKO mice on day 1 and day 2 after cerulein treatment (replicates n=3). Arrows indicate positive puncta signals, while dashed arrows denote diffuse cytoplasmic expression.

In sum, HBP1 ablation increases pro-inflammatory cytokine gene programs and acinar necrosis while impairing anti-inflammatory and cellular response gene programs, such as autophagy, upon injury.

### Loss of Hbp1 in AP impairs autophagic flux, which HBP1 directly regulates

To determine whether impaired autophagy gene programs resulted in the accumulation of autophagosomes, we investigated p62 and LC3 proteins in HBP1 CKO and WT pancreata after cerulein treatment. In HBP1 CKO mice, the number of LC3- and p62-positive puncta was increased, whereas in HBP1 WT mice, LC3 expression was diffusely distributed throughout the cytoplasm, indicating delayed autophagosome clearance in HBP1 CKO mice (Fig. 5E-L, Fig. S5).

To verify whether HBP1 activation was sufficient to regulate the autophagy gene program, we created inducible HBP1-expressing human normal pancreatic ductal epithelial cells (PDEC) and performed RNA-seq 24 hours post-HBP1 induction (Fig. S6A-C, Table S5). As expected, HBP1 overexpression downregulated Cyclin D genes while upregulating p21, BAD, FOXO1 and FOXO3 (Fig. S6D-F).

Surprisingly, HBP1 upregulation significantly enhanced the autophagy gene program (Fig. 4L-M, S6G). HBP1 overexpression functionally enhanced autophagic flux, as evidenced by decreased levels of the autophagy substrate p62, while the LC3-II/I ratio remained unchanged under basal conditions and upon inhibition of autophagosome-lysosome fusion with chloroquine (CQ) (Fig. 4N-O). These findings suggest that HBP1 upregulation promotes both autophagosome degradation and the autophagy-related gene program.

In sum, these findings suggest that impaired cellular protective mechanisms in HBP1 CKO pancreata contribute, at least in part, to severe pancreatitis driven by impaired autophagy.

### Loss of Hbp1 delays the onset of oncogenic KRAS-driven PanIN formation and PanIN progression

Since HBP1 is expressed in PanINs and associated with invasive tumors in pancreatitis (Fig. 1), and given that KRAS drives pancreatic neoplastic lesions, we investigated whether HBP1 influences the onset and progression of KRAS-driven PanINs. Thus, we generated *p48^Cre^; LSL-Kras^G12D^; Hbp1^flox/flox^* mice (HBP1-null or homozygous KCH), which allow both activating an oncogenic Kras and deleting Hbp1 in pancreatic epithelial cells (Fig. 6A). As an HBP1 WT control, littermate *Hbp1^+/flox^* mice (HBP1-competent or heterozygous KCH) were used. Both homozygous and heterozygous KCH mice were born at the expected frequency. Mouse body weight and normalized pancreatic weight did not differ between sexes or among genotypes (Fig. S7A-B, Table S6).

**Figure 6.**
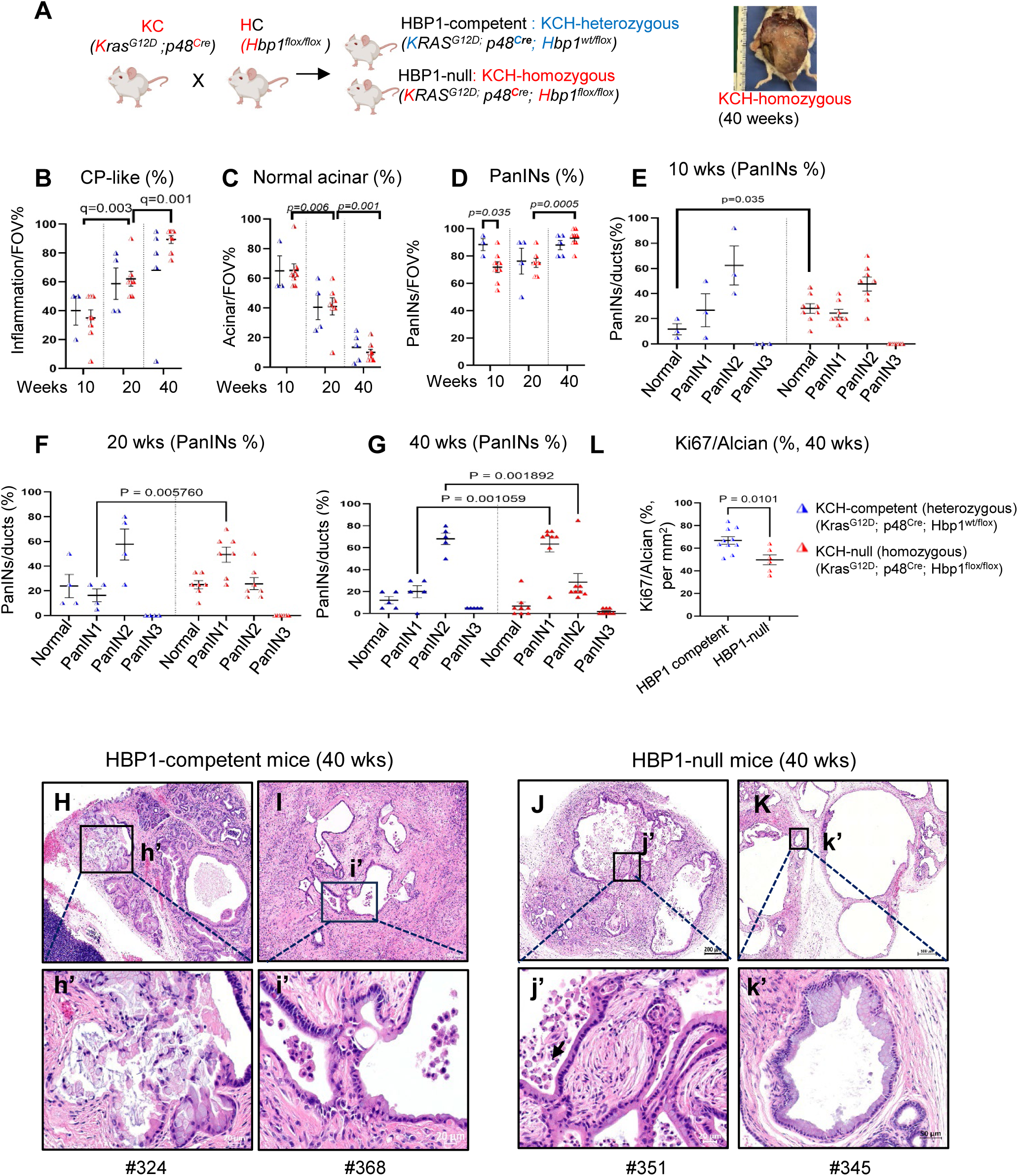
PanIN progression is delayed in HBP1 null mice (KCH-homozygous) compared to HBP1 competent mice (KCH-heterozygous). A. Diagram and representative image of KCH mice. B-D. The fraction of pancreatic parenchyma represents CP-like inflammation (B), acinar (C), and PanINs (D) The study included 24 KCH-null mice (n=8 per time point) and 12 KCH-competent mice (n=3 at 10 weeks, n=4 at 20 weeks, and n=5 at 40 weeks) (statistical significance, Welch t-test). The fraction of PanINs is significantly lower in KCH null mice compared to KCH competent mice at 10 weeks but significantly increases from 20 to 40 weeks in KCH null mice (D, Welch t-test). E-G. The fraction of total ducts representing each PanIN stage at 10 weeks (E), 20 weeks (F), and 40 weeks (G) of KCH-competent and null mice. The fraction of ducts representing PanIN1 and PanIN2 significantly differed between these two genotypes by 20 weeks and 40 weeks, respectively (Welch t-test). H-K. Representative images of the highest-grade PanIN lesions in HBP1-competent and null mice at 40 weeks. The ductal epithelium is thickened with papillary projections and exhibits nuclear atypia, including loss of basal nuclear polarity and pleomorphism, characteristic of PanIN2 to PanIN3 lesions (H, inset h’). The pancreatic ductal epithelium near regions of strong inflammation displays irregular glandular structures with large pleomorphic nuclei and luminal necrosis. It is surrounded by highly desmoplastic stromal cells (I, inset i’). HBP1-null mice developed high-grade PanIN3 lesions characterized by mild nuclear atypia and cytological atypia, including luminal necrosis (arrow in j’) and budding of small clusters of epithelial cells into the lumen (J, inset j’). Extensively dilated pancreatic cystic lesions with small low-grade PanIN1B/PanIN2 were frequently observed near CP-like inflamed areas (K, inset k’) (scale bars: 200, 500, and 20 µm). L. The ratio of Ki67 expressing cells over Alcian blue positive PanINs per mm^2^ at 40 weeks. (six random regions of each mouse are measured, HBP1-competent mice n=4, null-mice n=8, Welch t-test).

By ten weeks, both HBP1-competent and null mice developed CP-like inflammation with mixed PanINs affecting over half of the acinar pancreatic parenchyma (Fig. 6B-D). With age, most of the acinar pancreatic parenchyma was replaced by intense CP-like inflammation with diffuse PanINs in both groups (Fig. 6B-D). HBP1-null KCH mice exhibited delayed PanIN development at 10 weeks (Fig. 6D-E). While PanINs progressed to higher grades in both groups with age, HBP1-null mice showed a slower progression compared to HBP1-competent mice (Fig. 6E-G).

PanIN lesions in KCH mice were observed to originate from acinar cells via metaplasia and spontaneously from ductal cells by 10 weeks (Fig. S7C-D). By 40 weeks, both HBP1- competent and HBP1-null pancreata exhibited a range of PanINs, including high-grade PanIN3-like lesions with cytological atypia in a small fraction of pancreata (Fig. 6H-K, Table S6B). However, while PanIN3 lesions were present in all HBP1-competent mice, particularly near areas of severe inflammation (Fig. 6H-I), high-grade PanIN3 lesions were observed in only 3 out of 7 HBP1 CKO mice (Fig. 6J, Table S6B). In contrast, the majority of PanINs near inflamed areas in HBP1-null mice were low-grade PanIN1B lesions or cystic structures (Fig. 6K). Supporting this finding, the proportion of proliferating cells within Alcian Blue positive PanINs was significantly lower in HBP1-null mice than in HBP1 competent mice at 40 weeks (Fig. 6L, S7E-H). In contrast, the overall number of proliferating cells across the entire pancreas was not significantly different (Fig. S7I), suggesting that HBP1-null mice predominantly formed less aggressive PanINs.

In contrast, KC mice, regardless of littermate status, primarily harbored focal PanINs while maintaining a largely intact acinar compartment. They developed the full spectrum of PanINs, including PanIN3, by 20 weeks and progressed to an invasive phenotype by 40 weeks (Fig. S7J–N).

In summary, HBP1 ablation in KC mice delays the initiation of oncogenic KRAS-driven PanIN formation and restricts PanIN progression, potentially through mechanisms that remain to be fully elucidated.

## DISCUSSION

A deeper understanding of its molecular regulators is crucial to better predict the outcomes of AP and prevent its recurrence. Our study demonstrates that HBP1 expression increases in pancreatitis and is associated with disease progression. Furthermore, HBP1 ablation exacerbates acinar inflammatory damage through necrosis and autophagy dysregulation. In the presence of oncogenic *KRAS*, HBP1 ablation delayed the PanIN formation and slowed its progression to higher-grade lesions. These findings suggest that HBP1 is critical in preventing pancreatitis while also promoting PanIN progression, potentially through its role in maintaining cellular homeostasis.

Necrotizing pancreatitis, which has a worse prognosis than interstitial edematous pancreatitis, is marked by increased acinar cell loss^4, 5^. Disruptions in acinar homeostasis and impaired autophagy contribute to disease severity by exacerbating cell death^6–8^. Our data show that HBP1 ablation leads to acinar cell dropout through necrosis rather than apoptosis, accompanied by dysregulated autophagy in HBP1 CKO pancreata. Specifically, HBP1 ablation decreases key autophagic gene expression and results in autophagosome accumulation, while HBP1 overexpression enhances autophagic flux and autophagic gene programs, including LAMP3 and TFEB, which are also implicated in pancreatitis development in both mice and humans^42, 43^.

mTORC1 suppresses both the early initiation and late degradation phases of autophagic flux by inhibiting ULK1 and TFEB, two key regulators of autophagy^44, 45^. Notably, mTORC1 signaling was significantly downregulated in iHBP1-OE PDEC cells (Fig. S6H), suggesting that HBP1 may enhance autophagy, at least in part, by inhibiting mTORC1. This aligns with prior findings that HBP1-null mice exhibit increased mTORC1 activation^46^. Together, these findings indicate that HBP1 regulates autophagosome formation and promotes autophagosome-lysosome degradation, serving as a protective gatekeeper against acinar necrosis upon injury. The precise mechanisms by which HBP1 regulates overall autophagic flux remain to be determined.

While dysregulated autophagy and acinar cell death in HBP1 CKO mice likely exacerbate inflammation, HBP1 may also function upstream of immune-modulatory pathways to limit pancreatic inflammation and promote recovery. Macrophage recruitment plays a crucial role in pancreatitis severity^47^, and our data show that HBP1 CKO pancreata exhibit increased expression of proinflammatory cytokine genes, which are essential for monocyte and macrophage infiltration. Notably, HBP1 overexpression activated type I IFN pathways (Fig. S6I), whereas HBP1 CKO pancreata exhibited significant downregulation of Reg family genes, which protect against pancreatic damage by modulating immune responses^33, 34^. HBP1 binding sites are significantly enriched in the regulatory regions of immune response genes^23^. Thus, we cannot exclude the possibility that HBP1 counteracts inflammation by directly transactivating anti-inflammatory pathways, which warrants further investigation.

While HBP1 protects against pancreatitis-related injury and is dispensable for injury-induced metaplasia, its ablation has the opposite effect in oncogenic KRAS-driven pancreatic neoplasia. HBP1-null KCH mice exhibit delayed PanIN formation and a lower incidence of high-grade PanINs compared to HBP1-competent KCH mice. Since PanIN progression is closely linked to autophagy, which supports the survival of both PanINs and PDAC^48–50^, further studies are needed to determine whether HBP1 promotes PanIN progression through autophagy-dependent mechanisms.

In conclusion, upregulated HBP1 in AP limits inflammatory injury by maintaining acinar homeostasis and reducing macrophage infiltration. However, in the presence of an oncogenic KRAS mutation, HBP1 facilitates PanIN progression. Targeting both HBP1 and autophagy presents a potential dual therapeutic strategy: enhancing autophagy in pancreatitis to resolve inflammation and prevent recurrence while inhibiting autophagy in KRAS-driven lesions to block PanIN progression. This dual-targeting approach may offer a precision medicine strategy for effectively managing pancreatitis and preventing its recurrence and progression.

## Abbreviations

AP: Acute pancreatitis
CP: Chronic pancreatitis
FNA: Fine-Needle Aspiration
PDAC: Pancreatic ductal adenocarcinoma
PanIN: Pancreatic intraepithelial Neoplasia
IPMN: Intraductal Papillary Mucinous Neoplasm
IPMC: Intraductal papillary mucinous carcinoma
ADM: Acinar-to-ductal metaplasia
HBP1: HMG-box transcription factor 1
CCK: Cholecystokinin
TF: Transcription factor
IF: Immunofluorescence
IHC: immunohistochemistry
H&E: Hematoxylin & Eosin
TMA: Tissue microarray
qRT-PCR: Quantitative reverse transcription-PCR
CKO: Conditional knockout
ROC: Receiver operating characteristic
AUC: Area Under Curve
Reg: Regenerating gene
UMAP: Uniform Manifold Approximation and Projection for Dimension Reduction
DEG: Differentially regulated genes

## Author contributions

**Conceptualization**: JK; **Data curation**: SL, TE, DG, SY, JL, and TM; **Formal Analysis**: SL, TE, ECM, SY, JL, MB, DG, MS, JLM, KC, SPW, TM, and JK; **Investigation**: SL, TE, ECM, SY, JL, MB, DG, MS, SH, JLM, KC, SPW, TM, and JK; **Methodology**: SL, TE, ECM, MB, DG, JL, MS, COM, VMS, JLM, KC; **Resource**: VMS, DK, JLM, RCS; **Writing**: SL and JK; **Supervision**: KC, RCS, and JK; **Funding Acquisition**, JK.

## Data Transparency Statement

All data and analytic methods will be made available to other researchers. Spatial Transcriptomic data and RNA-sequencing will be available on NCBI GEO. Additional results are available in supplementary figures and tables as resource datasets and any requests will be fulfilled by the lead contact, Jungsun Kim.

## ACKNOWLEDGMENTS

We thank Drs. Cote and Zimmers for comments on the manuscript; Drs. Ohtsuka and Kitano (Kyoto) and Fedorov (OHSU Transgenic Mouse Models) for fertilized the embryos of Hbp1^floxed/floxed^ strain and generating p48^Cre^; HBP1^floxed/floxed^ mice, respectively; K. Rice, M. Davenport, and Y. Wang for their technical assistance; OHSU Shared Resource (Histopathology, Transgenic Mouse Models); Brenden-Colson Center for Pancreatic Care, BioLibrary, and the Oregon Pancreas Tissue Registry (IRB #3609). Life Science Editors edited this manuscript. Graphical abstracts were created using Biorender.com.

## SUPPLEMENTARY FIGURES

### Supplementary figure legends

**Figure S1.**
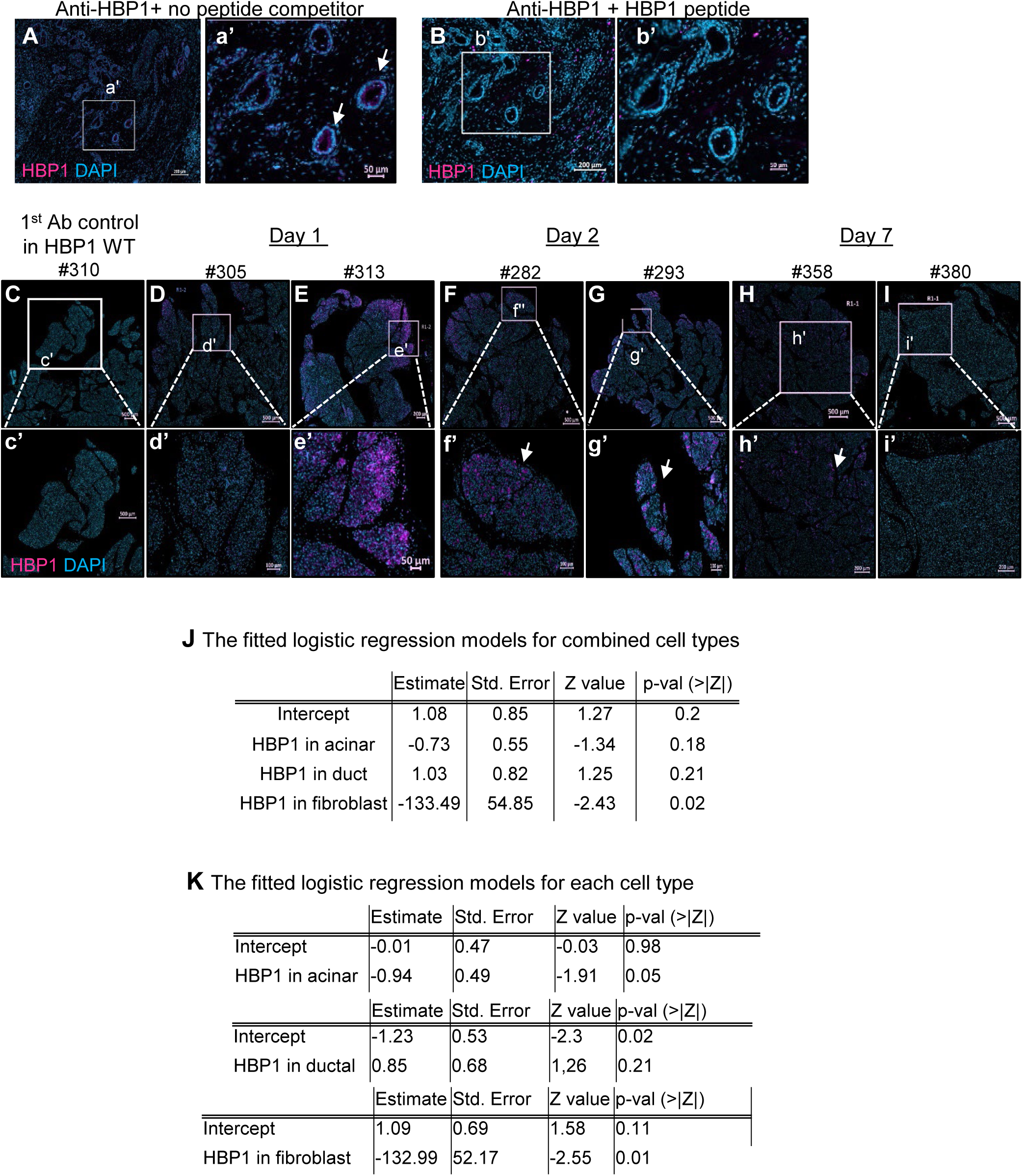
Pancreatic HBP1 accumulation gradually decreases throughout tissue regeneration. Related to Figure 1. A-B. Validation of HBP1 antibody specificity by peptide competition assay. A rabbit polyclonal anti-HBP1 antibody was incubated with no competitor (A) or with a 100-fold molar excess of recombinant HBP1 peptide (B) for 30 min at room temperature and then applied to the immune staining. The excess HBP1 competes for the signal. The arrows indicate HBP1 staining. C-I. Expression of HBP1 during cerulein-treated pancreatitis. Images from no primary antibody control (C) and two additional mice are shown in D-I. The arrows indicate HBP1 staining. J-K. The fitted logistic regression model for combined cell types (J) and each cell type (K).

**Figure S2.**
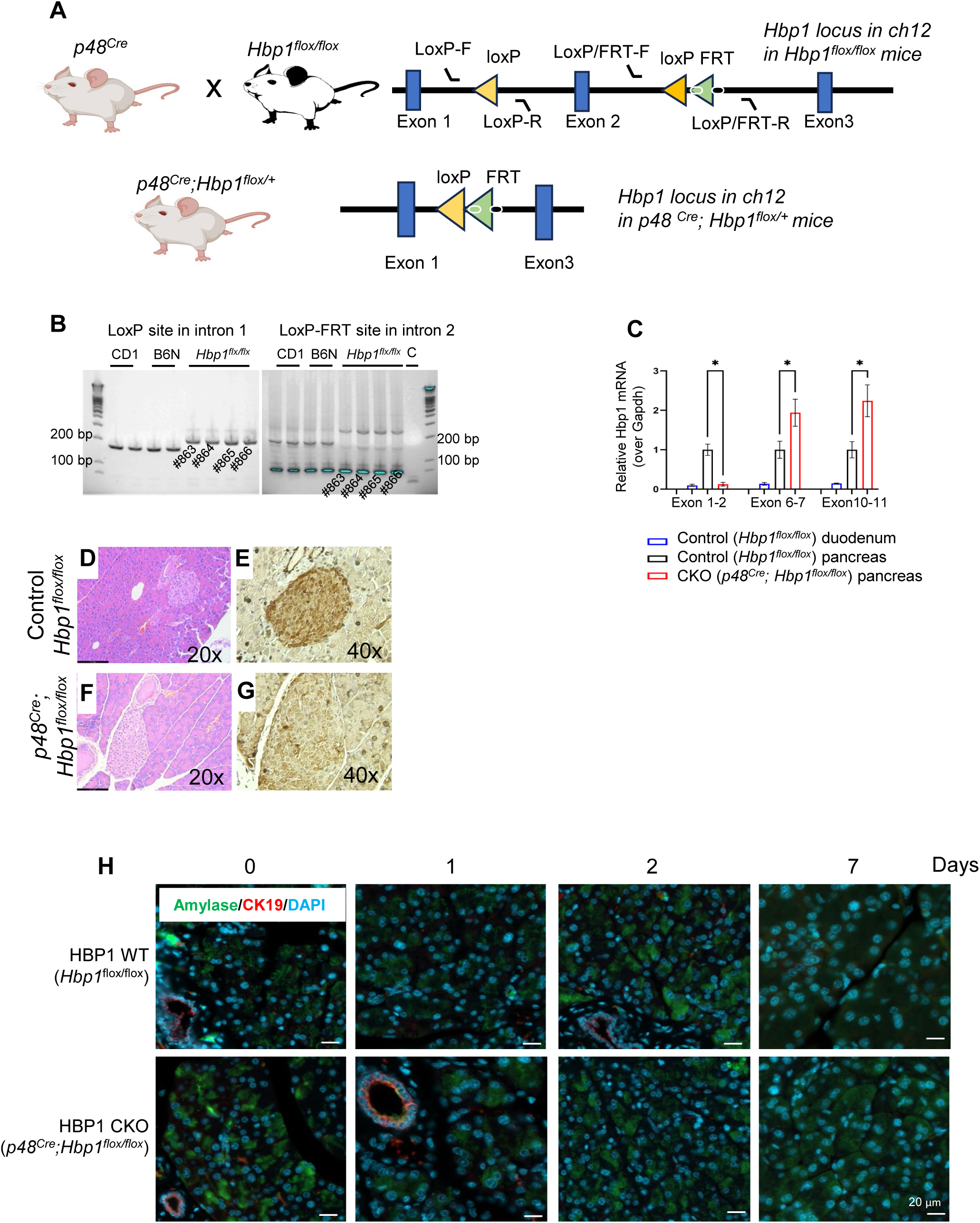
Generation of pancreatic Hbp1 CKO mice. Relevant to Figure 2. A. A schematic diagram of a part of the *Hbp1* locus (chr12:31920507-31956193) of *Hbp1^flox/flox^* mice. One loxP site is integrated into the intron 1 and another loxP and Frt sites are integrated into the intron 2. After crossing with *p48^Cre^* mice, cre recombinase deletes the DNA between the two loxP sites in *Hbp1^flox/flox^* mice and generates *p48^Cre^; Hbp1^flox/WT^*. To make homozygotic *p48^Cre^; Hbp1^flox/flox^*, the heterozygous mice are repetitively bred with each other. B. Representative agarose gel images to verify the insertion of loxP and loxP/FRT sites in intron 1 and intron 2, respectively. Using Hbp1 loxP site primers in intron1, Hbp1 WT and floxed alleles give rise to 173 bp and 219 bp, respectively. Using Hbp1loxP FRT site primers in intron 2, Hbp1 WT and floxed alleles produce 184 bp and 272 bp, respectively. C. Confirmation of missing the part of Hbp1 mRNA encoded by the exon 2 using real-time PCR. Real-time PCR primers are designed to detect parts of the *Hbp1* mRNAs encoded by exon1/2, exon6/7, and exon 10-11. Duodenum and pancreatic tissues from control mice with intact *Hbp1* alleles are used as negative and positive controls, respectively. The Y-axis shows *Hbp1* mRNA levels relative to *Gapdh.* The pancreata from CKO mice lacking *Hbp1* alleles show the complete absence of *Hbp1* exon1/2 mRNAs, which is a level relatively similar to negative controls, compared to control mice with intact *Hbp1* alleles (* p<0.005, Welch t-test, n=6 technical replicates). D-G. Confirmation of the absence of HBP1 protein in HBP1 CKO mice. Representative H&E (D, F), HBP1 IHC (E, G) on pancreatic islet cells from control and CKO mice. H. Amylase and CK19 staining on pancreata from HBP1 WT control (upper) and HBP1 CKO (lower) mice on day 0, 1, 2, 7 after cerulein treatment (n=3).

**Figure S3.**
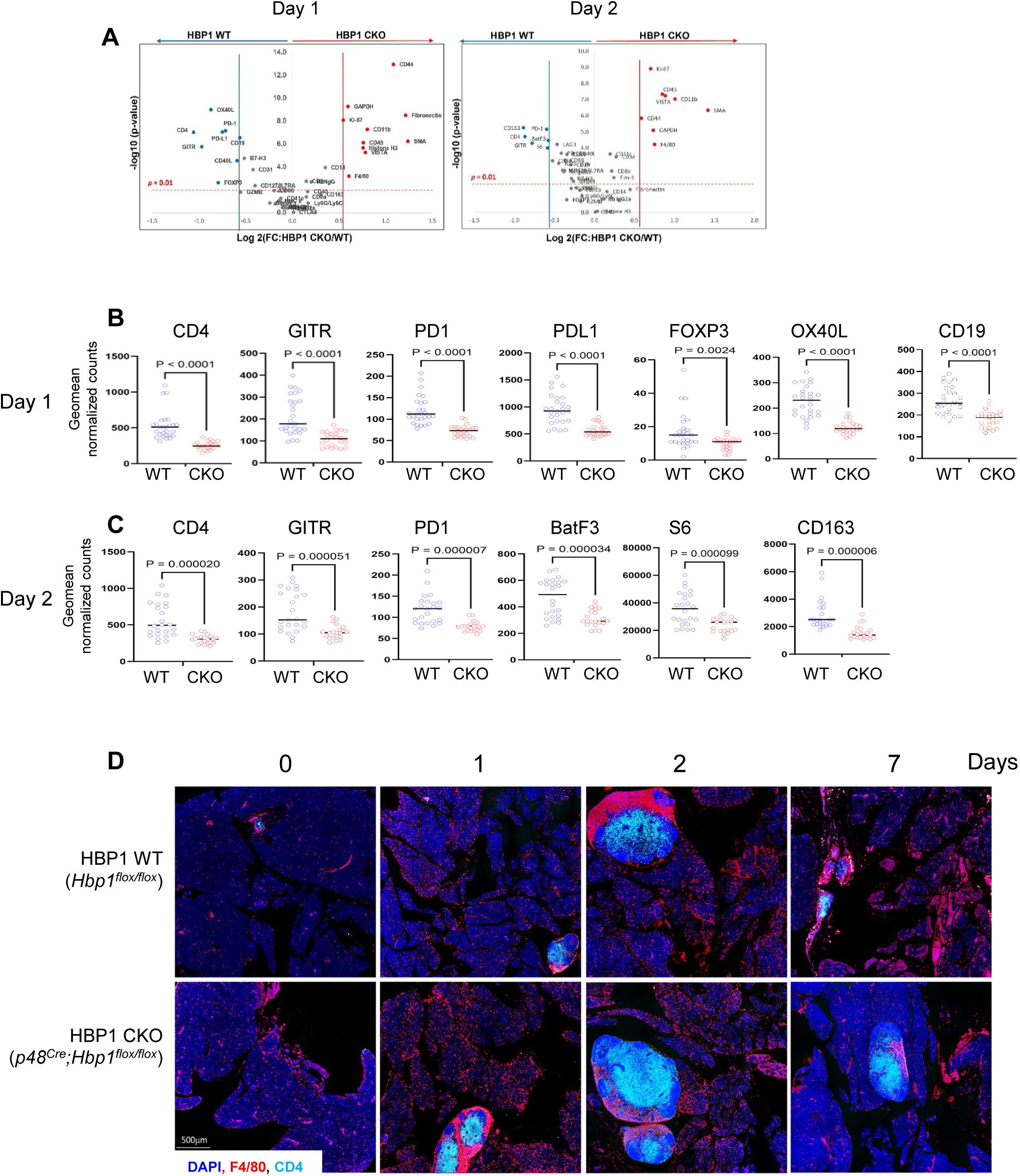
Relevant to Figure 3. A. Volcano plots of differentially expressed immune markers with geomean normalized counts. The X-axis indicates log 2-fold differences between CKO and WT pancreata, and the Y-axis indicates -log 10 (p-value). We selected proteins that significantly increased (Red) or decreased (Blue) over 1.5-fold in CKO compared to WT (p<0.01). Non-significantly regulated proteins are in gray. Raw N-count was normalized with signal-to-noise ratio (SNR). Significantly regulated proteins are defined based on p-value and fold changes (FC: HBP1 CKO/WT). P-values are less than 0.01, and the expression ratio of markers in CKO over WT is larger than 1.5-fold. Log2 (FC)>0.585. B-C. Top proteins downregulated in HBP1 CKO pancreata day 1 (B) and 2 (C) post-cerulein treatment. D. The representative images of F4/80^+^ and CD4^+^ T cells in the peripancreatic lymphoid node tissues during injury after cerulein treatment.

**Figure S4.**
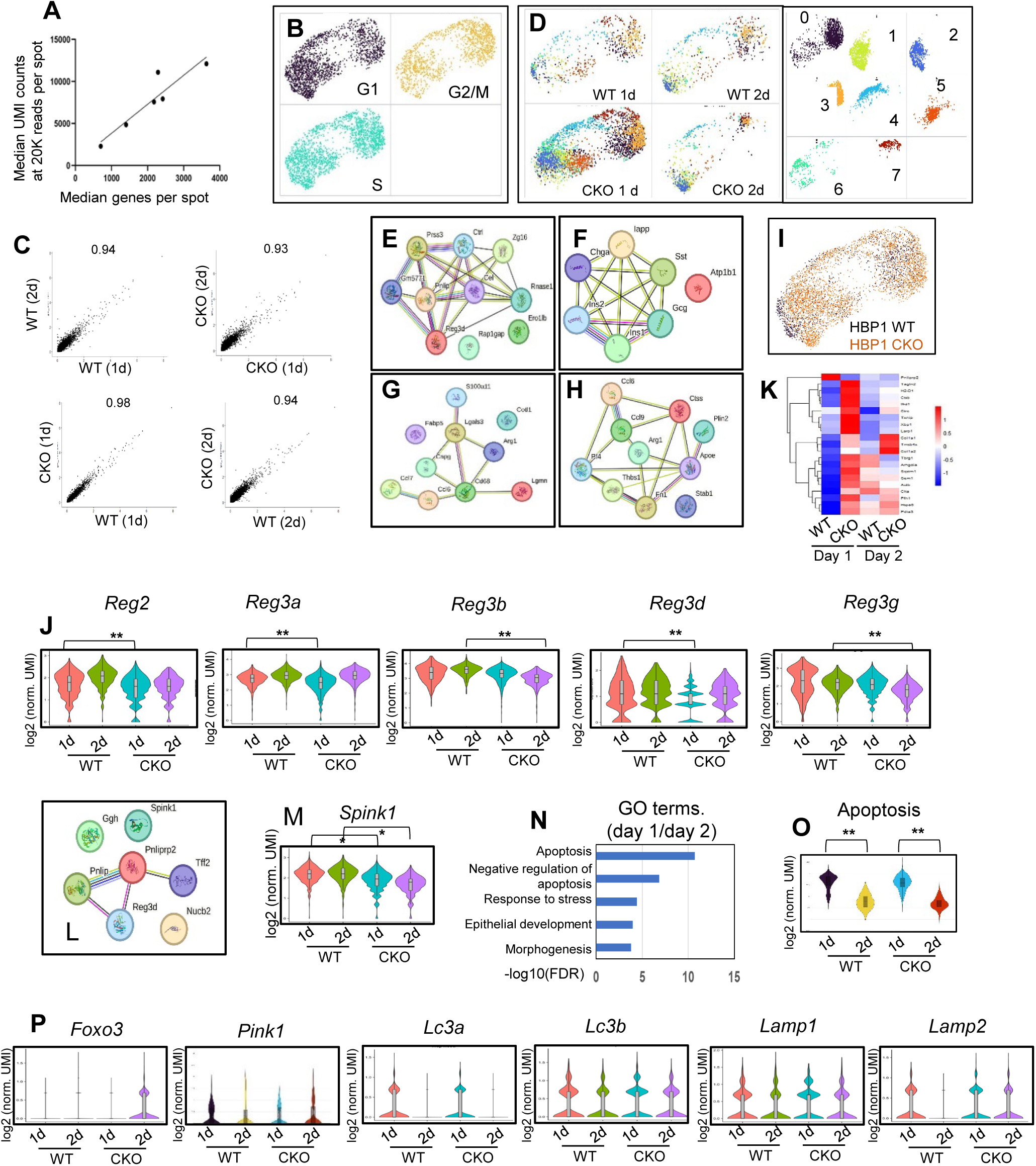
Relevant to Figure 4. A. Scatter plot of median gene numbers and median UMI counts at 20K reads in each spot. B. UMAP of cell cycle genes. C. Spearman correlation coefficient among samples. D. UMAPI of eight clusters in each group. E-H. Network analysis of top 10 DEG in group 2 (E), group 6 (F), group 0 (G), and group 3 (H). I. UMAP of aggregated clusters in each group. J. Expressions of the Reg family genes *Reg2, Reg3a, Reg3b, Reg3d* and *Reg3g*. K. Heatmap of differentially expressed genes in group 2 cells between CKO and WT at day 1. The Heatmap key indicates the z-score. L. Network of differentially downregulated genes in CKO compared to WT among group 2. M. The expression of the Spink1 gene (* FDR<0.05). N. GO terms of genes that are increased on day 1 compared to day 2. O. The expression of apoptotic pathway genes (* FDR<0.05). P. Expression of autophagy genes (y-axis is average log 2 normalized UMI).

**Figure S5.**
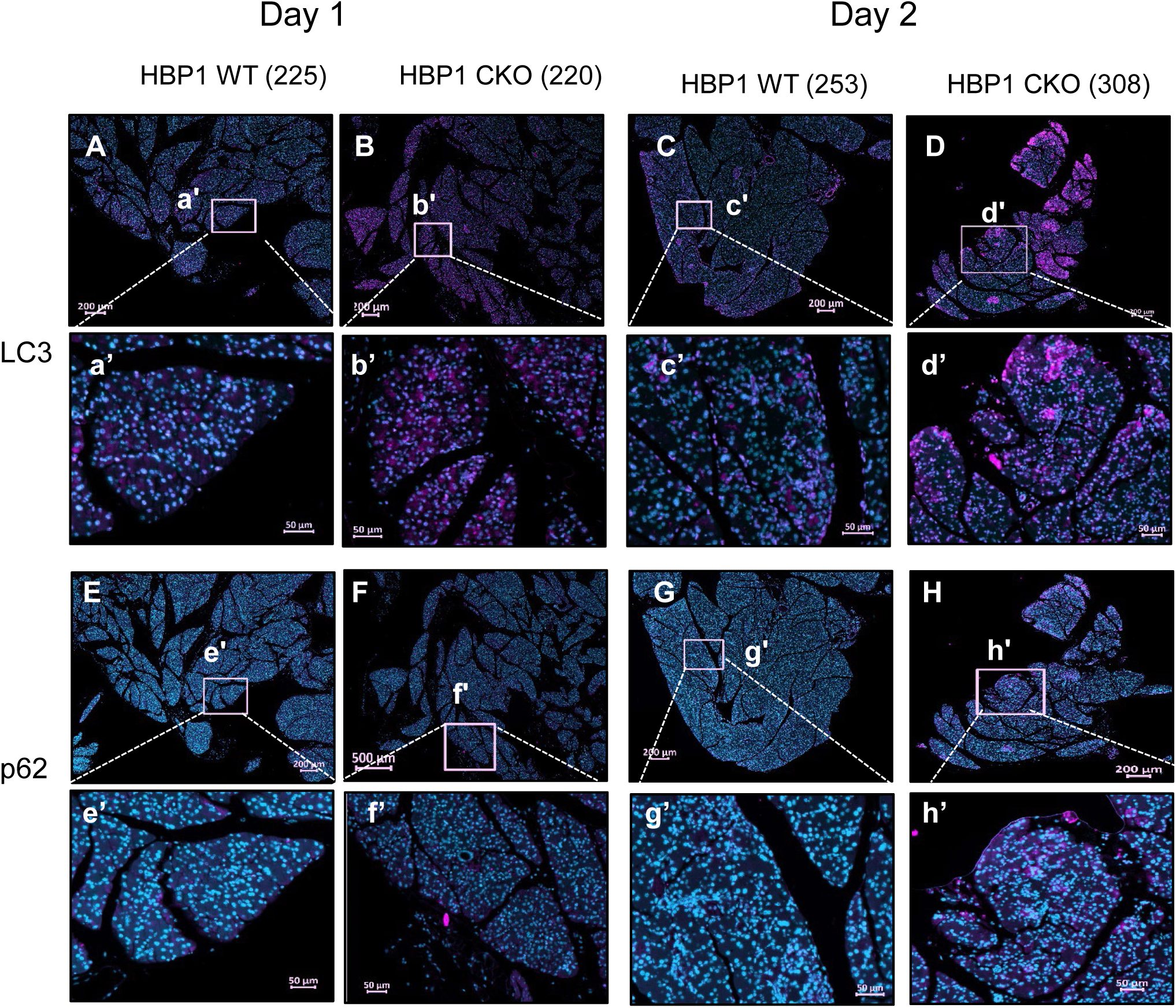
Relevant to Figure 5. Additional immune staining of LC3 (A-D) and p62 (E-H) autophagosome marker proteins in pancreata from HBP1 WT and CKO mice on day 1 and day 2 after cerulein treatment (LC3 or p62 stained in pink, DAPI stained in blue).

**Figure S6.**
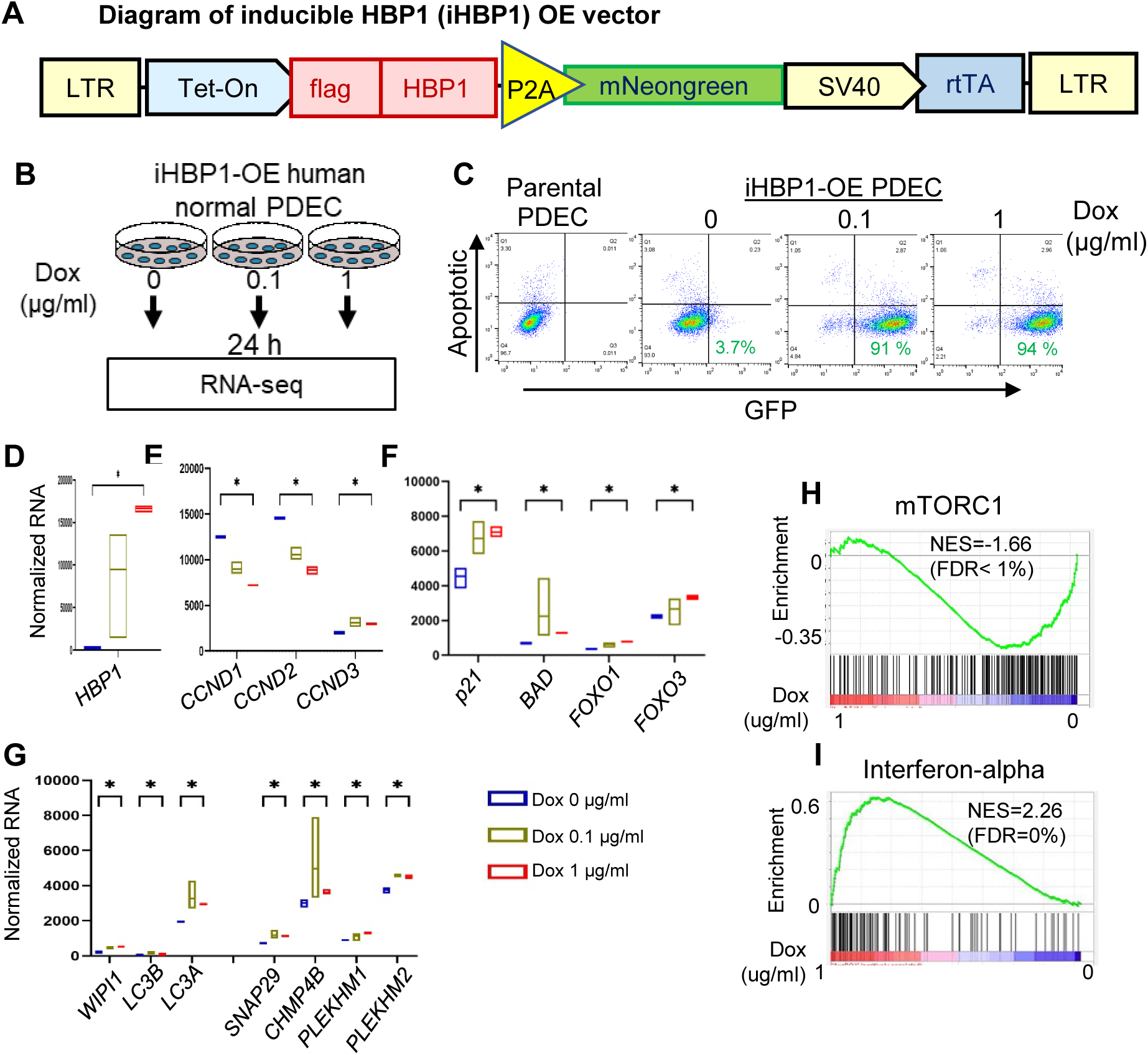
Relevant to Figure 4. Confirmation of autophagy gene program regulated by HBP1 in human normal pancreatic ductal epithelial cells (PDEC)^33^. A. Diagram of inducible human HBP1 (iHBP1) overexpressing (OE) lentiviral vector for generating inducible HBP1 expressing PDEC. B. Schematic flowchart of RNA-seq from iHBP1-PDEC (n=3). C. Validation of iHBP1-PDEC by flow cytometry. D-F. Validation of doxycycline-dependent expression of HBP1 (D) and HBP1-regulated cyclin D genes (E) and senescence (p21, FOXO1, FOXO3), and apoptosis (BAD) (F). The X-axis indicates concentrations of doxycycline (0, 0.1, 1 µg/ml), and the Y-axis indicates normalized DESEQ2 counts (*FDR<0.05). G. Expression of genes in autophagy pathways among DEG (*FDR<0.05). H-I. GSEA for genes that are decreased (H) and increased (I) in HBP1 overexpressing PDEC.

**Figure S7.**
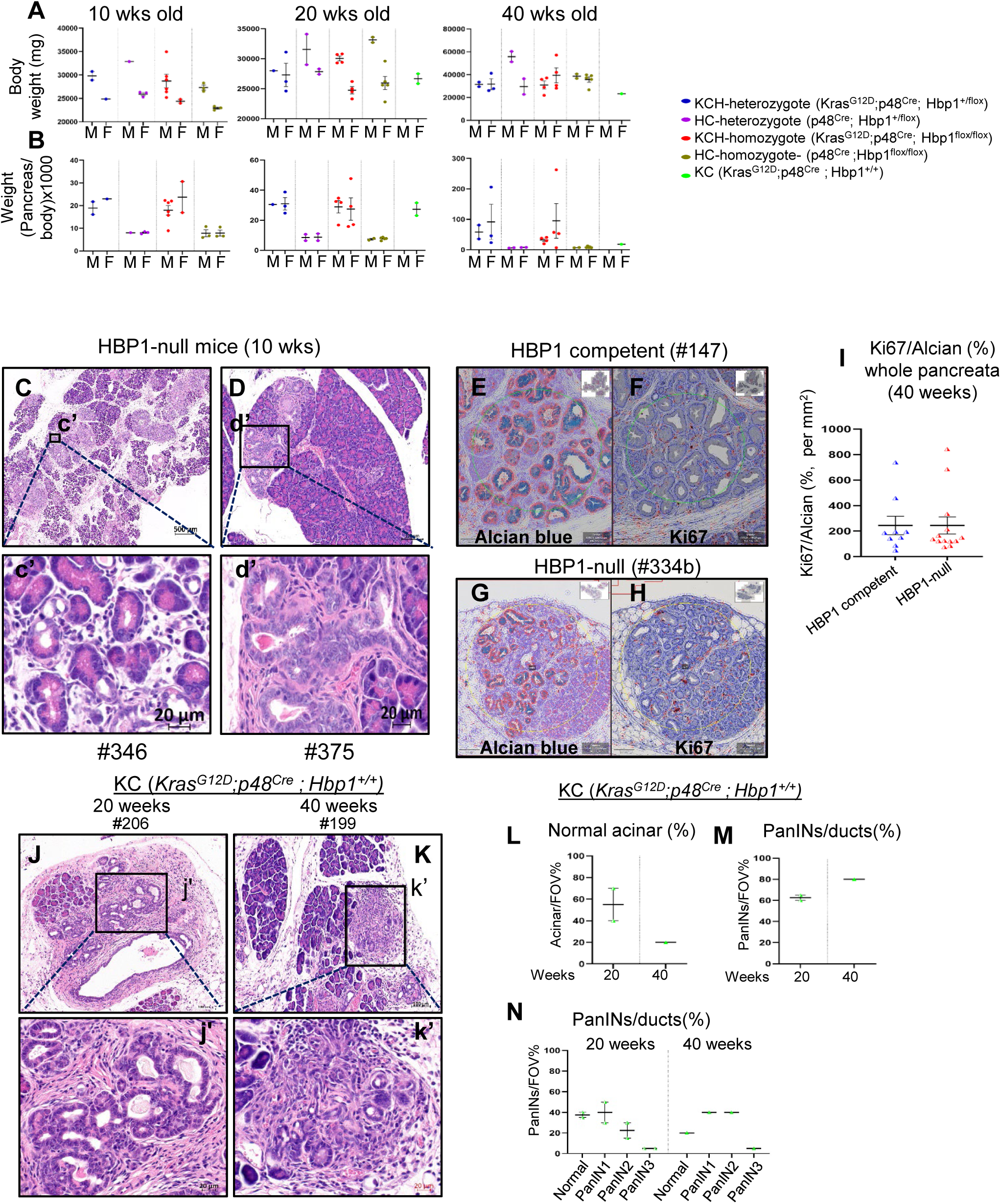
Relevant to Figure 6 KCH mice. A-B. Body weight (A) and pancreatic weight normalized by body weight (B) in each sex for KC and KCH mice. There was no difference between sex. C-D. Representative H&E images of HBP1-null KCH pancreata at 10 weeks. By 10 weeks of age, HBP1-null mice generate PanIN-like cells from acinar origins (C, inset c’) and spontaneously from ductal cells (d, inset d’). E-H. Representative images of quantification of Ki67 (F, H) within Alcian blue (E, G) using QuPath. I. Quantification of Ki67 in the entire pancreata (40 weeks). J-K. Representative H&E images of the highest grade of KC mice at 20 (J, inset j’) and 40 (K, inset k’) weeks. Focal PanIN3-like lesions near normal acinar cells exhibit irregular glandular structures with nuclear pleomorphism (J, inset j’). By 40 weeks, KC mice developed invasive PDAC, characterized by small, highly irregular, and distorted ductal structures with poorly defined organization, nuclear atypia, and pleomorphism (K, inset k’). L-N. The fraction of normal acinar (L), PanINs (M), and ducts representing each PanIN in 20 and 40 weeks (N) in KC mice. F-G.

## SUPPLEMENTARY TABLES

**Supplementary Table S1. Primer Sequences.**

**Supplementary Table S2. The information about pancreatitis TMA core.**

Table S2A. Summary of TMA core samples

Table S2B. Demographics of the patients participating.

Table S2C. The fraction of tissues expressing different levels of HBP1 in pancreatitis TMA cores

Table S2D. Statistical analysis of HBP1 expression levels in pancreatitis TMA cores

**Supplementary Table S3. The information about mice that have been used for the cerulein-induced AP model.**

Table S3A. The body weight and pancreatic weight information of mice were used for the study.

Table S3B. Statistical analysis of sex.

Table S3C. Statistical analysis of body weight, pancreatic weight, and normalized pancreatic weight by body weight.

**Supplementary Table S4. The information about visium spatial transcriptomics.**

Table S4A. Median genes and UMI counts per spots

Table S4B. Gene ontology of group 0 (macrophage/neutrophils)

Table S4C. Gene ontology of group 2 (pancreatic acinar)

Table S4D. Gene ontology of group 3 (myofibroblast, myeloid)

Table S4E. List of genes significantly upregulated in group 6 (pancreatic islet)

**Supplementary Table S5. The GSEA analysis on iHBP1-OE-PDEC with and without doxycycline induction.**

Table S5A. GSEA for Dox 1 ug/ml treated iHBP1-OE-PDEC.

Table S5B. GSEA for Dox 0 ug/ml iHBP1-OE-PDEC control

**Supplementary Table S6. The information about KCH mice.**

Table S6A. The body weight and normalized pancreatic weight of KC and KCH mice.

Table S6B. The histology analysis of KC and KCH mice.

Table S6C. The quantification of Alcian Blue and Ki67 positive cells in KCH mice (40 weeks).

Table S6D-1. The ratio of Ki67/Alcian blue (per nm2) in total samples.

Table S6D-2. The ratio of Ki67/Alcian blue (per nm2) in only PanIN lesions without cystic lesions.

## What You Need to Know

### BACKGROUND AND CONTEXT

Pancreatitis is an inflammatory disease of the pancreas and a risk factor for pancreatic cancer. HBP1, a transcription factor, is linked to pancreatic cancer progression, but its role in pancreatitis remains unclear.

### NEW FINDINGS

HBP1 protects against severe pancreatic inflammation but also promotes early cancerous changes in the presence of an oncogenic mutation, highlighting its dual role in disease progression.

### LIMITATIONS

The significant negative association of HBP1 level and the presence of tumors in fibroblasts highlights a potential protective mechanism, while trends in acinar and ductal cells suggest the need for further investigation. Future studies with larger cohorts are necessary to confirm these findings and explore the underlying mechanisms.

### CLINICAL RESEARCH RELEVANCE

Understanding HBP1’s role in acute pancreatitis and early pancreatic cancer development could inform new therapeutic strategies to prevent disease recurrence and progression, ultimately improving patient outcomes.

### BASIC RESEARCH RELEVANCE

This study identifies HBP1 as a key regulator of pancreatic inflammation and autophagy, linking pancreatitis severity to early cancerous changes. These findings provide new insights into the molecular mechanisms driving pancreatitis progression and its transition to pancreatic cancer, offering potential therapeutic targets.

### LAY SUMMARY

HBP1 protects the pancreas from severe inflammation but also promotes early cancerous changes when a mutation is present. Targeting HBP1 may help manage pancreatitis and reduce pancreatic cancer risk.

